# Timing anti-PD-L1 checkpoint blockade immunotherapy to enhance tumor irradiation

**DOI:** 10.1101/2024.08.06.606845

**Authors:** Steve Seung-Young Lee, Joanna Pagacz, Sera Averbek, David Scholten, Yue Liu, Stephen J. Kron

## Abstract

**Background:** The ability of radiotherapy (RT) to drive anti-tumor immunity is limited by adaptive resistance. While RT induces inflammation and recruits activated tumor-infiltrating lymphocytes (TILs) including cytotoxic T lymphocytes (CTLs), the resulting radiation- and IFNγ-dependent PD-L1 expression restores an immunosuppressed tumor microenvironment. Unleashing an effective anti-tumor response may require precise sequencing of RT and checkpoint blockade immunotherapy (CBI) to block PD-L1 signaling before it can mediate its suppressive effects.

**Methods:** Flank tumors formed in BALB/c mice with syngeneic CT26 colon or 4T1 mammary carcinoma cells were treated with otherwise ineffective doses of ionizing radiation (10 Gy) followed by CBI (0.2 mg anti-PD-L1, i.v.) after 0, 1, 3, 5 or 7 days, comparing tumor response. Anti-PD-L1 delivery was measured by fluorescence, TILs by flow cytometry and immunofluorescence, PD-L1 expression by immunohistochemistry, and tumor size by calipers. **Results** In both CT26 and 4T1 tumors, 10 Gy alone resulted in a transient growth delay associated with infiltrating CTLs peaking at 3 days and PD-L1 at 5 days. CTLs returned toward baseline by 7 days, consistent with adaptive resistance. Anti-PD-L1 failed to potentiate radiation except when injected 5 days after 10 Gy, which prevented CTL depletion and led to tumor elimination. Potentially contributing to compound effects, anti-PD-L1 penetrated tumors and bound PD-L1 more efficiently after irradiation.

**Conclusions:** Optimal timing to exploit radiation-induced permeability to enhance CBI delivery and interrupt adaptive resistance by blocking PD-L1 as it peaks may offer a general strategy to enhance external beam radiotherapy by protecting activated TILs and potentiating anti-tumor immune response.

## BACKGROUND

Binding of programmed cell death protein 1 receptor PD-1 (CD279), expressed by cytotoxic T lymphocytes (CTLs) and other immune effectors, to its ligand PD-L1 (B7-H1, CD274), expressed by myeloid and cancer cells in the tumor microenvironment (TME), limits CTL expansion, activation, cytotoxicity, and survival, driving immune evasion.^1,2^ Multiple checkpoint blockade immunotherapy (CBI) antibodies are now approved that disrupt PD-1/PD-L1 signaling to restore CTL function in PD-L1-expressing tumors.^3,4^ A complementary use for CBI may be in combination with external beam radiotherapy (RT^5,6^) and other treatments^7,8^ that induce PD-L1, toward overcoming adaptive resistance. While RT promotes immunogenic cell death (ICD) and release of proinflammatory signals,^9,10^ its vaccine-like effects are typically self-limiting, attributed to activated T cells releasing IFNγ in the tumor and inducing PD-L1 expression to drive rebound immunosuppression.^11,12^ In some preclinical models, PD-1/PD-L1 blockade can potentiate anti-tumor immune response following RT, but translation to the clinic has been challenging.^13,14^

Along with cytotoxic and immune effects, RT also transiently disrupts the tumor microvascular permeability barrier,^15,16^ facilitating extravasation of circulating macromolecules including therapeutic antibodies. Radiation-induced permeability peaks within several days, associated with damage to pericytes and disruption of the basal lamina.^17^ Notably, infiltration of inflammatory cells displays similar kinetics after irradiation, suggesting common mechanisms.

Here, we reexamined anti-PD-L1 antibody as a means to potentiate radiation’s vaccine-like effects. Consistent with IFNγ-dependent adaptive resistance after radiation, CTL infiltration was followed by increased PD-L1 expression and then CTL depletion. A single dose of anti-PD-L1 at the peak of PD-L1 expression protected the CTL infiltrate and potentiated tumor elimination. These data point to a tight window of opportunity to disrupt adaptive resistance and release an effective anti-tumor immune response after radiotherapy.

## Materials and methods

### Cell lines and animal models

4T1 and CT26 cells (ATCC) were cultured in RPMI-1640 media with 10% FBS (Denville), 2 mM L-glutamine, and 100 units/mL penicillin/streptomycin. Negative test results for mycoplasma and a murine virus panel were confirmed for 4T1 and CT26 cell samples by IDEXX RADIL.

Animal studies were approved by the University of Chicago IACUC. Tumors were formed subcutaneously on the right flank of female BALB/c (6-8 weeks old, Envigo) or NSG mice with 5×10^5^ 4T1 or 2.5×10^5^ CT26 cells. Tumors were treated after 14 days with 0.2 mg anti-PD-L1 (10F.9G2, BioXCell) in 0.1 mL PBS (pH 7.4) injected through the tail vein or with 10 Gy (IR, 2.5 Gy/min, X-RAD 225Cx, Precision X-Ray) and then anti-PD-L1 after 1, 3, 5, or 7 days. Tumors were measured by calipers along the major (a) and minor (b) axes and volume calculated as (a×b^2^)/2. We randomly assigned tumor mice to the groups and treated them in a blind manner.

### In vitro inhibitor treatment

5×10^4^ CT26 cells were seeded into 6 well plate and allowed to adhere overnight. The cells were treated with 1 μM of STING inhibitor C-178 (Selleckchem) or 1 uM of Ruxolitinib (Selleckchem) 2 h before radiation. Cells were irradiated with 10 Gy (^60^Co, 7.09 cGy/sec, GammaCell, Nordion) and protein was extracted 3 days after radiation.

### Immunodetection

For flow cytometry, tumors excised 2 days after treatment were digested using a Mouse Tumor Dissociation Kit (Miltenyi) and cells strained through a 70 µm filter into RPMI-1640 medium were pelleted by centrifugation and resuspended in 0.1 mL PBS. After 15 min on ice with FcR block (1.0 μg, BioLegend), cells were stained with anti-CD3-Alexa 700 or Pacific Blue (17A2), anti-CD8a-BB515 (53-6.7), anti-CD4-PE/Cy7 (GK1.5), anti-CD45-Pacific Blue or Alexa 700 (30-F11), and anti-CD49b-APC (DX5) fluorescent conjugates (Biolegend) for 45 min at 4°C. Cells washed with PBS and stained with 10 µg/mL propidium iodide (PI) were analyzed along with costained CompBeads on a Fortessa cytometer (BD) and data analyzed by FlowJo (TreeStar) and Prism (GraphPad).

For immunohistochemistry, excised tumors were fixed, embedded, sectioned, processed and stained on a Bond RX with the PDL-1C protocol using anti-PD-L1 (E1L3N, Cell Signaling) and Bond Polymer Refine Detection (Leica). Tissue sections were imaged at 40× on a Pannoramic SCAN BF (Perkin Elmer). PD-L1 staining was quantified with ImageJ (version 1.53h) at six randomly selected regions to obtain mean intensity ± SEM.

For immunofluorescence, sections were manually processed and stained with anti-PD-L1 (17952-1-AP, Proteintech, 1:600), anti-CD45 (10-F11, BD, 1:750) or anti-CD8a (4SM15, eBioscience, 1:500) and anti-perforin (E3W4I, Cell Signaling, 1:700), each overnight at 4°C, washed in TBS-Tween 20 (0.05%), stained with anti-rabbit-DyLight594 (Jackson), anti-rat-647 (Jackson) and DAPI (1 µg/ml), mounted in Prolong Gold (Invitrogen) and imaged at 40x using an SP5 confocal microscope (Leica).

For Western blotting, CT26 cells treated with 10 Gy were analyzed at 1, 3, 5 or 7 days with anti-PD-L1 (ab213480, Abcam, 1:1000) and anti-rabbit-HRP (Cytiva, 1:10000) using anti-β-actin-HRP as loading control.

### Tracking antibody delivery

Anti-PD-L1 (10F.9G2, BioXCell) and anti-CD31 (MEC13.3, Biolegend) diluted in PBS pH 8.0 were labeled with NHS-activated DyLight 488, 594, or 633 (Thermo) overnight at 4°C, dialyzed against PBS pH 7.4 at 4°C for 3 days, and stored at 4°C.

At 5 days after 4T1 and CT26 tumors were irradiated with 10 Gy, mice were injected *i.v.* with anti-PD-L1-DyLight 594 (0.2 mg in 0.1 mL PBS) and tumors excised after 30 min. Tumors were embedded in OCT (Tissue-Tek), frozen at -80°C and sectioned at -20°C. Sections dried at room temperature were rehydrated in PBS, stained with anti-PD-L1-DyLight 488 and anti-CD31-DyLight 633, washed and imaged with an SP8 confocal microscope (Leica).

### Quantification and statistical analysis

Data were analyzed using Prism (GraphPad) and represented as mean ± SEM. Comparison of two groups was by Mann-Whitney U test. Repeated-measure two-way ANOVA (mixed-model) followed by the Bonferroni post hoc test was used for analysis of tumor growth curves. p≤0.05 was considered statistically significant.

## Results

### Anti-PD-L1 potentiates radiation effects

In prior studies,^18^ we observed that circulating 10F.9G2 anti-PD-L1 peaks within minutes after tail vein injection and then returns nearly to baseline by 24 h, offering a means to examine optimal sequencing with IR. Thus, on day 0, we injected 4T1 or CT26 carcinoma cells into the right hindlimb of female BALB/c mice. Treating tumors on day 14 with a single *i.v.* injection of 0.2 mg anti-PD-L1 antibody did not affect tumor growth kinetics while a single 10 Gy dose of ionizing radiation (IR) induced a transient growth delay (figure 1A, online supplemental figure 1A). To examine combination therapy, tumors were treated with 10 Gy at 14 days and then with anti-PD-L1 at 1, 3, 5, or 7 days after IR (figure 1B-C, online supplemental figure 1B-C). A prolonged growth delay compared to 10 Gy alone was observed only when anti-PD-L1 was administered 5 days post-IR (p<0.05, figure 1C-E, online supplemental figure 1C-E). For 4T1 and CT26 tumors formed in immunodeficient NSG mice, 10 Gy and anti-PD-L1 antibody were ineffective alone or combined (online supplemental figure 2), confirming a role for lymphocytes in the growth delay.

**Figure 1.**
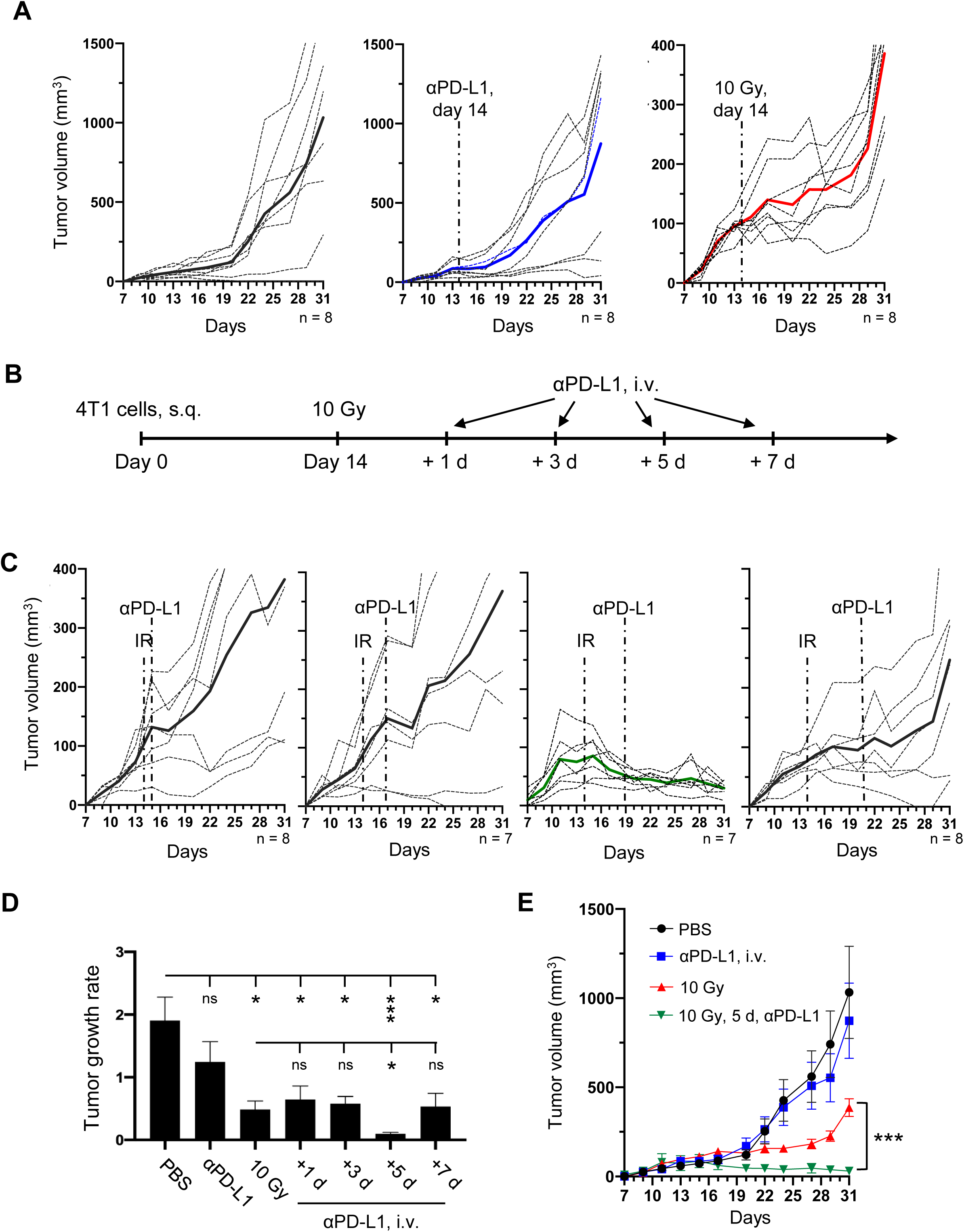
PD-L1 antibody at 5 days after radiation leads to tumor suppression. (**A**) 4T1 tumors were established in BALB/c immunocompetent mice at day 0 and treated at day 14 with PBS, anti-PD-L1, or 10 Gy, showing individual tumor growth profiles and mean. (**B**) Schema for combination therapy where tumors irradiated at day 14 are treated with anti-PD-L1 after 1, 3, 5, or 7 days. (**C**) Growth profiles for individual tumors and mean for combination treatment, with timing as indicated. (**D**) Mean tumor growth rate calculated as tumor size on day 7 and day 29. Mean tumor growth rate ± SEM. (**E**) Mean tumor size for each group comparing controls to optimal combination therapy. *** p<0.01, *p≤0.05, n.s. p>0.05 n=7-8.

### PD-L1 expression and TILs in the response to radiation and combination therapy

To investigate the effect of IR on PD-L1 expression, we collected 4T1 and CT26 tumors at 1, 3, 5 or 7 days after 10 Gy and performed anti-PD-L1 immunohistochemistry (figure 2A-C). IR increased PD-L1 expression in tumors in a time-dependent manner in both models, reaching a maximum at 5 days after IR (4T1 p=0.022, CT26 p=0.022, figure 2B, C).

**Figure 2.**
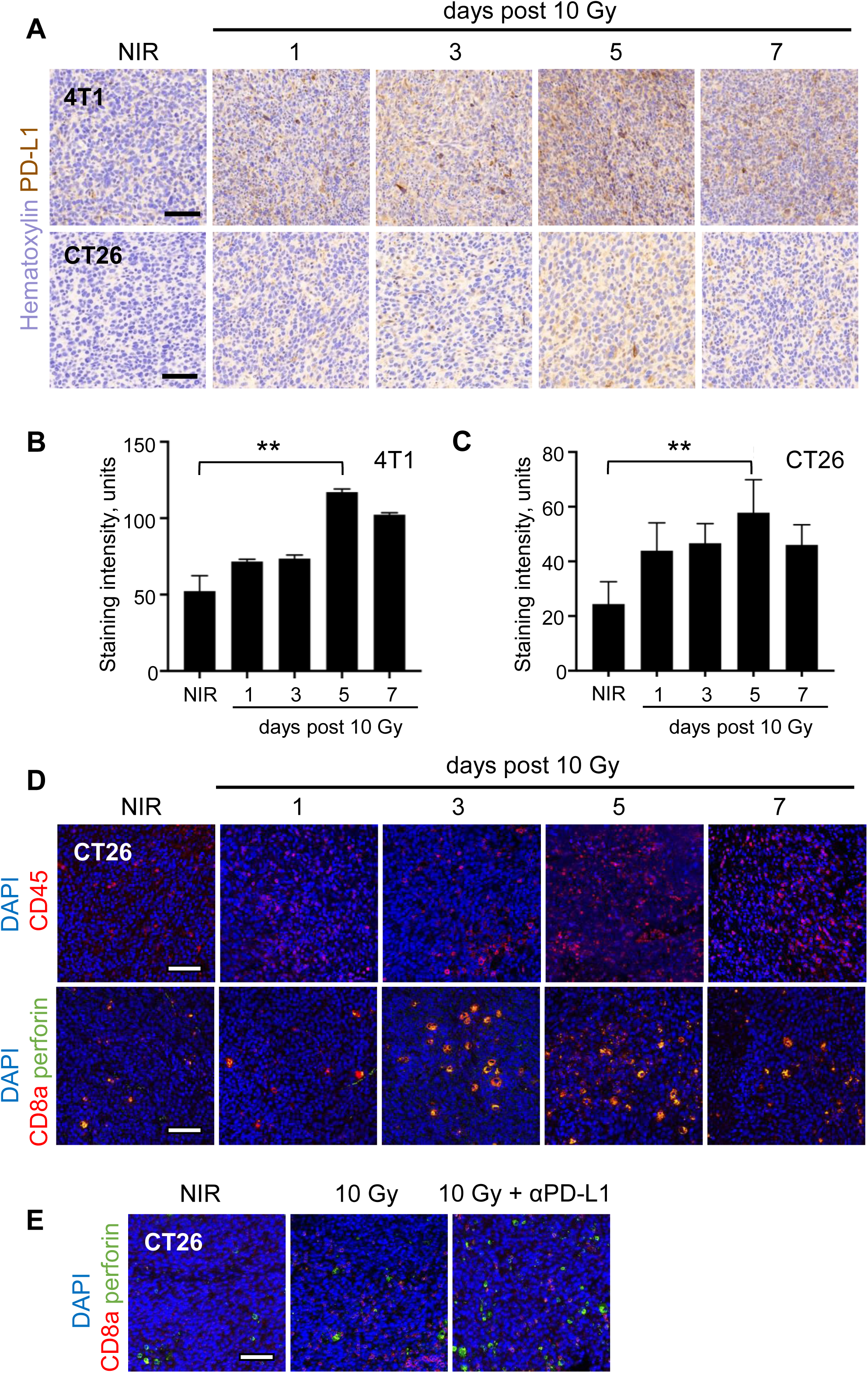
PD-L1 expression and TIL accumulation after radiation. (**A**) PD-L1 immunohistochemistry in 4T1 and CT26 tumors treated with 10 Gy and harvested after 1, 3, 5, or 7 days. Relative quantification of PD-L1 expression in (**B**) 4T1 and (**C**) CT26 tumors. Mean ± SEM. (**D**) Tumor sections were stained with anti-CD45 (red) or anti-CD8a (red) and anti-perforin (green) and DAPI (blue). (**E**) Tumors treated with 10 Gy alone or with anti-PD-L1 at 5 days and harvested at 7 days and stained with anti-CD8a (red) and anti-perforin (green). ** p<0.01. Scale bars: 50 µm

To examine CTL accumulation after radiation, CT26 tumors were excised at 1, 3, 5 or 7 days after 10 Gy, fixed, sectioned and stained with anti-CD45, CD8a and perforin antibodies. Tumors displayed heterogeneous staining, particularly at the edge of necrotic regions and the tumor capsule. Within the parenchyma, unirradiated tumors displayed a sparse, unactivated CTL infiltrate. By 3 days, CD45^+^ cells appeared to increase (figure 2D) and many of the CD8a^+^ TILs stained as perforin^+^. At 5 days, while CD45^+^ infiltrate further increased, intact CD8a^+^ cells decreased. At 7 days, the CD45^+^ infiltrate remained high, but the density of CD8a^+^ cells further decreased. While treatment with anti-PD-L1 at day 5 maintained CD8a and perforin at 7 days, CD8a^-^ perforin^+^ cells also increased (figure 2E), potentially representing NK cells and/or plasmacytoid DCs.^19,20^

To examine the immune response to combination treatment, 4T1 and CT26 tumors were excised 7 days after 10 Gy, with or without anti-PD-L1 on day 5. Tumors (n≥3 per group) were dissociated and the cell suspension immunostained for flow cytometric analysis of TILs (online supplemental figures 3 and 4). Anti-PD-L1 substantially increased CD45^+^ cells compared to radiation alone (4T1 p=0.11, CT26 p=0.032) with several TIL subsets appearing increased, including CD45^+^ CD3^+^ T cells, the CD4^+^ and CD8a^+^ T cell subsets and CD49b^+^ natural killer (NK) cells. CT26 tumors contained higher TILs than 4T1 tumors, perhaps reflecting higher immunogenicity, but responses were similar.

### Radiation facilitates anti-PD-L1 delivery

To track antibody delivery, mice were injected with anti-PD-L1-DyLight 594 and tumors were excised after 30 min. Then, sections were stained with anti-CD31-DyLight 633 to identify the microvascular endothelium and anti-PD-L1-DyLight 488 to label PD-L1 not reached by anti-PD-L1-DyLight 594 (figure 3). For both 4T1 (**A**) or CT26 (**B**), anti-PD-L1-DyLight 594 staining was limited to CD31^+^ perivascular regions in unirradiated tumors, leaving most of the tumor PD-L1 unbound and detectable with anti-PD-L1-DyLight 488. When injected 5 days after 10 Gy, anti-PD-L1-DyLight 594 displayed greater extravasation and deeper penetration, masking PD-L1 from detection by anti-PD-L1-DyLight 488.

**Figure 3.**
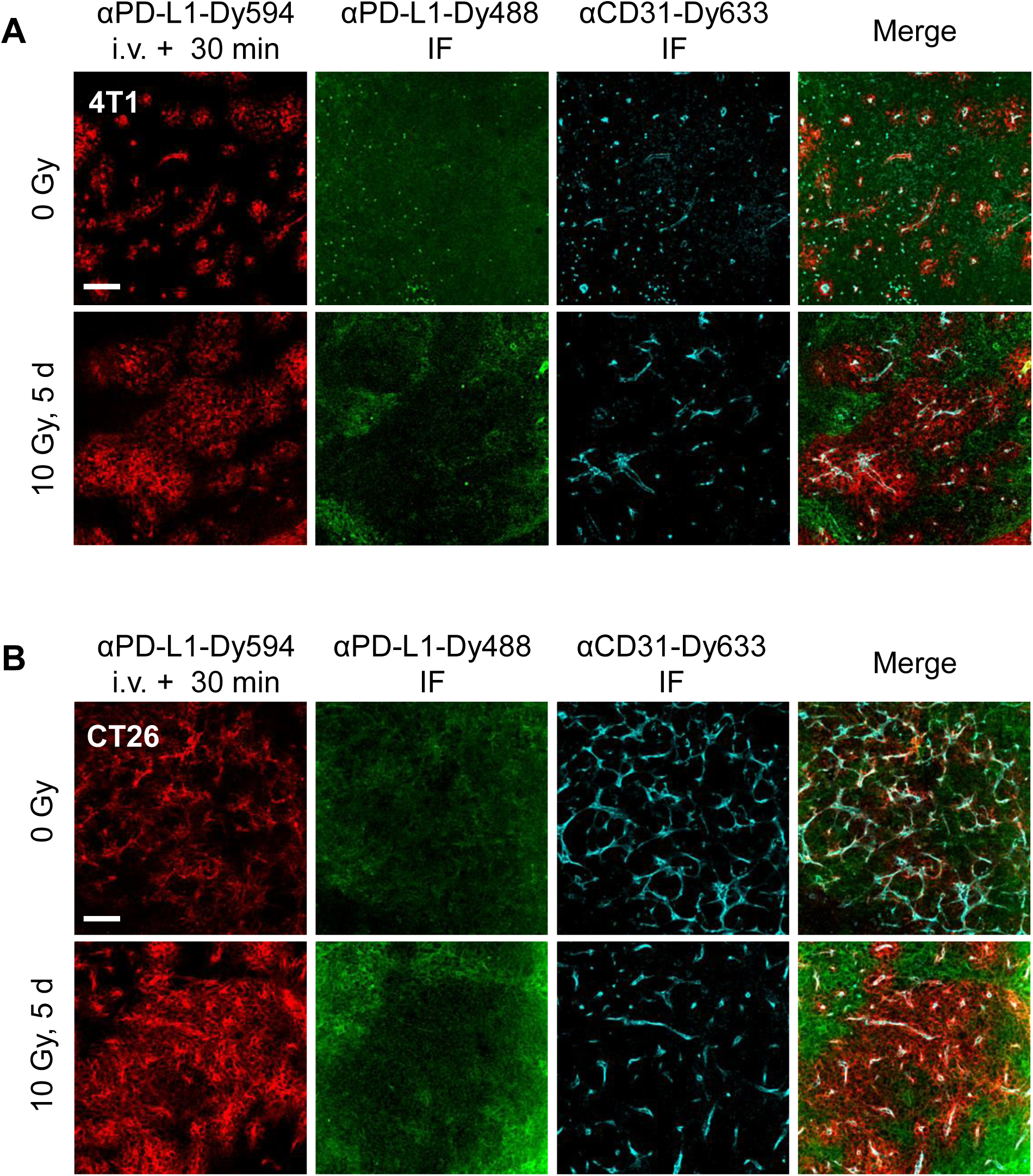
Radiation-induced permeability enhances anti-PD-L1 delivery. Distribution patterns of anti-PD-L1-DyLight 594 (red) in non-irradiated (upper row) or irradiated (lower row) of 4T1 (**A**) and CT26 (**B**) tumors. Staining with anti-PD-L1-DyLight 488 (green) indicates PD-L1 not bound by injected anti-PD-L1 and anti-CD31 indicates vascular endothelium (cyan). Scale bars: 100 µm.

### TILs determine PD-L1 expression after radiation

To evaluate cell-intrinsic effects on PD-L1 expression, CT26 cells were treated with 0 or 10 Gy, incubated for up to 7 days, and lysates analyzed by Western blot. PD-L1 was not induced until day 3, suggesting an indirect and/or delayed effect of radiation. Treatment with 10 Gy in the presence of STING or JAK inhibitors blocked PD-L1 induction at 3 days (figure 4B), suggesting a role for autocrine signaling. Co-culture of CT26 cells with mouse lymph node cells activated with anti-CD3/CD28 beads as a source of cytokines rapidly induced PD-L1 (figure 4B). To extend this comparison *in vivo*, CT26 tumors formed for 14 days in BALB/c or immunodeficient NSG mice were treated with 10 Gy and tumors excised after 5 days for analysis of PD-L1 expression (figure 4C). While PD-L1 was induced in NSG mice, this was markedly less than in BALB/c mice, implicating both cell-intrinsic signaling and effects of infiltrating, activated TILs in PD-L1 upregulation *in vivo*.

**Figure 4.**
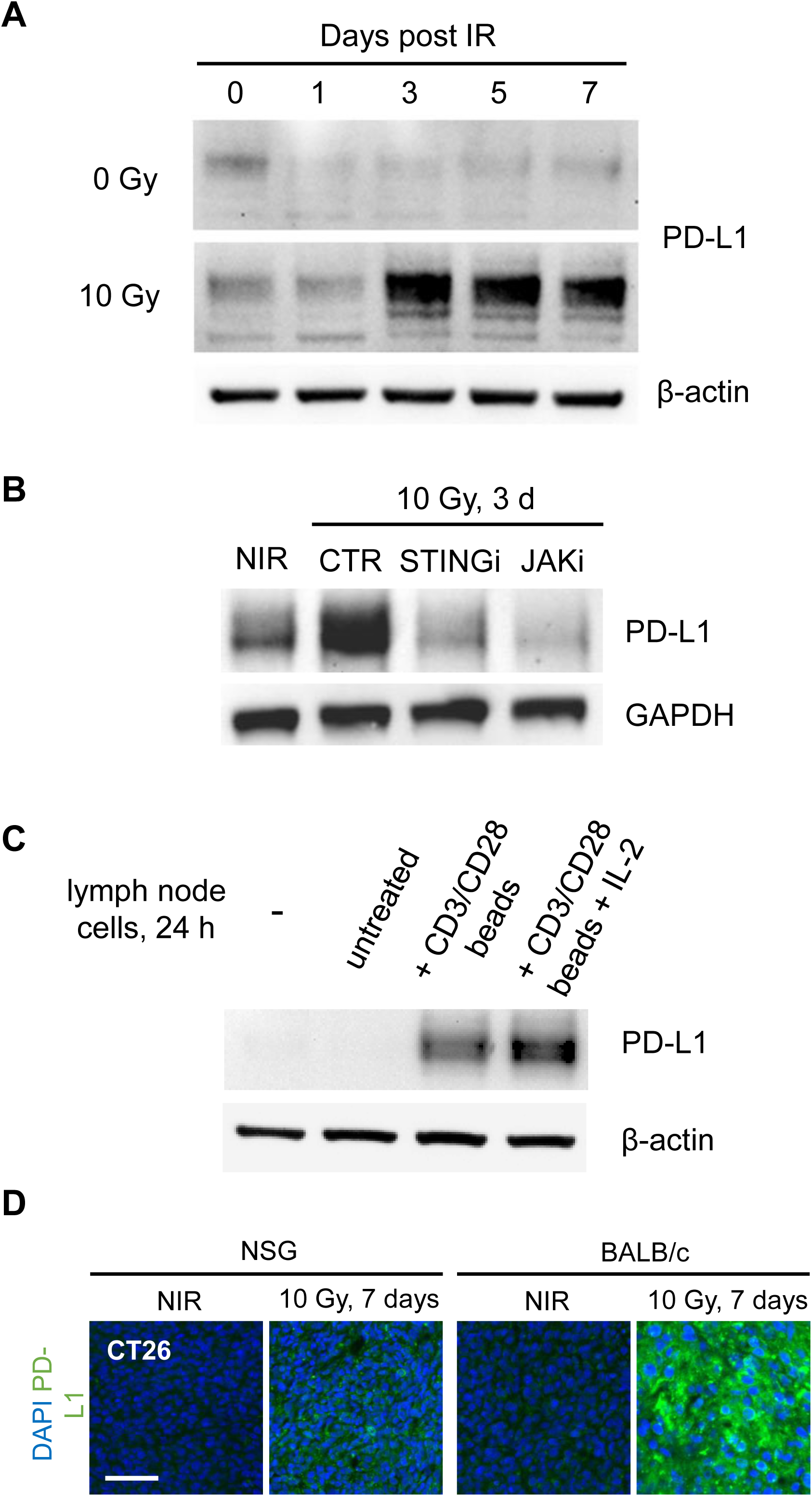
Intrinsic vs. paracrine PD-L1 upregulation in response to radiation. (**A**) CT26 cells were treated with 0 or 10 Gy and incubated for the indicated time and then cell lysates examined by anti-PD-L1 immunoblot. (**B**) CT26 cells were treated with 1 μM STING or JAK inhibitor prior to 0 or 10 Gy irradiation and cell lysates isolated after 3 days for anti-PD-L1 immunoblot. (**C**) CT26 cells were treated with the indicated agents or coincubated for 24 h with mouse lymph node cells, untreated, activated with anti-CD3/CD28 beads, or activated with anti-CD3/CD28 beads with IL-2 and lysates isolated for anti-PD-L1 immunoblot. (**D**) CT26 tumors in BALB/c or NSG mice were treated with a single 10 Gy dose, isolated after 7 days and stained with anti-PD-L1 antibody and imaged by immunofluorescence. Scale bar: 100 µm.

## Discussion

The anti-tumor immune response induced by radiation is typically short-lived as tolerance is rapidly restored via rebound immunosuppression.^21^ Among factors that induce PD-L1 in the tumor microenvironment after radiation, the most potent may be IFNγ released by activated TILs themselves.^11,12^ Overcoming this adaptive resistance may release a sustained anti-tumor immune response following radiation therapy, leading to tumor elimination.^22^

We found a relatively short interval after irradiation during which targeting PD-1/PD-L1 signaling was most effective in disrupting adaptive resistance. Most combinations of single doses of IR and anti-PD-L1 displayed only additive effects. However, injecting anti-PD-L1 at the peak of PD-L1 expression following irradiation resulted in marked growth inhibition and tumor elimination. Several factors may contribute to the specific timing but PD-L1 expression appeared to depend on the inflammatory infiltrate. PD-L1 induction followed TIL accumulation in BALB/c mice and was blunted in NSG mice deficient in T, B and NK cells. In turn, by disrupting the negative feedback loop, anti-PD-L1 antibody increased TILs and facilitated tumor elimination. Given this simple model, circulating or tissue markers of adaptive resistance could be used to guide timing of CBI after irradiation.

Another factor that may determine optimal timing of anti-PD-L1 effects is radiation-induced permeability which, like PD-L1 accumulation, develops over several days after irradiation.^17^ Notably, the transient loss of pericyte coverage and disruption of extracellular matrix after radiation may not only enhance macromolecular delivery but also facilitate leukocyte extravasation, potentially increasing the impact of properly timed CBI.

In summary, we describe a strategy of sequencing single doses of ionizing radiation and immune checkpoint blockade therapy to disrupt adaptive resistance and achieve strong anti-tumor response in multiple tumor models. We anticipate that clinical applications of similar treatment regimens may yield greater benefits to more patients.

## Supplemental figure legends

**Supplemental Figure 1.**
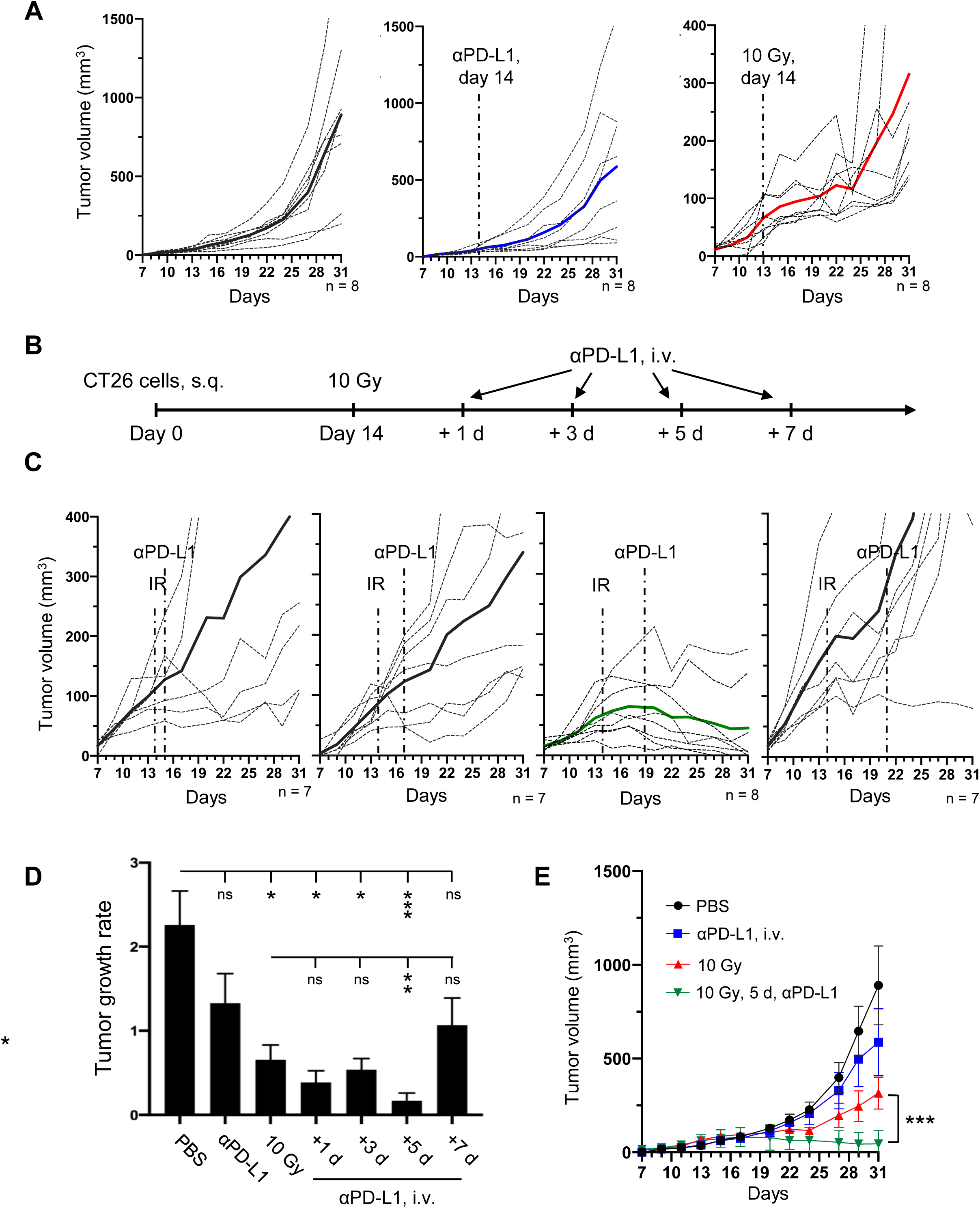
PD-L1 antibody at 5 days after radiation leads to tumor suppression. (**A**) CT26 tumors were established in BALB/c immunocompetent mice at day 0 and treated at day 14 with PBS, anti-PD-L1, or 10 Gy, showing individual tumor growth profiles and mean. (**B**) Schema for combination therapy where tumors irradiated at day 14 are treated with anti-PD-L1 after 1, 3, 5, or 7 days. (**C**) Growth profiles for individual tumors and mean for combination treatment, with timing as indicated. (**D**) Mean tumor growth rate calculated as tumor size on day 7 and day 29. Mean tumor growth rate ± SEM. (**E**) Mean tumor size for each group comparing controls to optimal combination therapy. *** p<0.01, * p≤0.05, ns p>0.05, n=7-8.

**Supplemental Figure 2.**
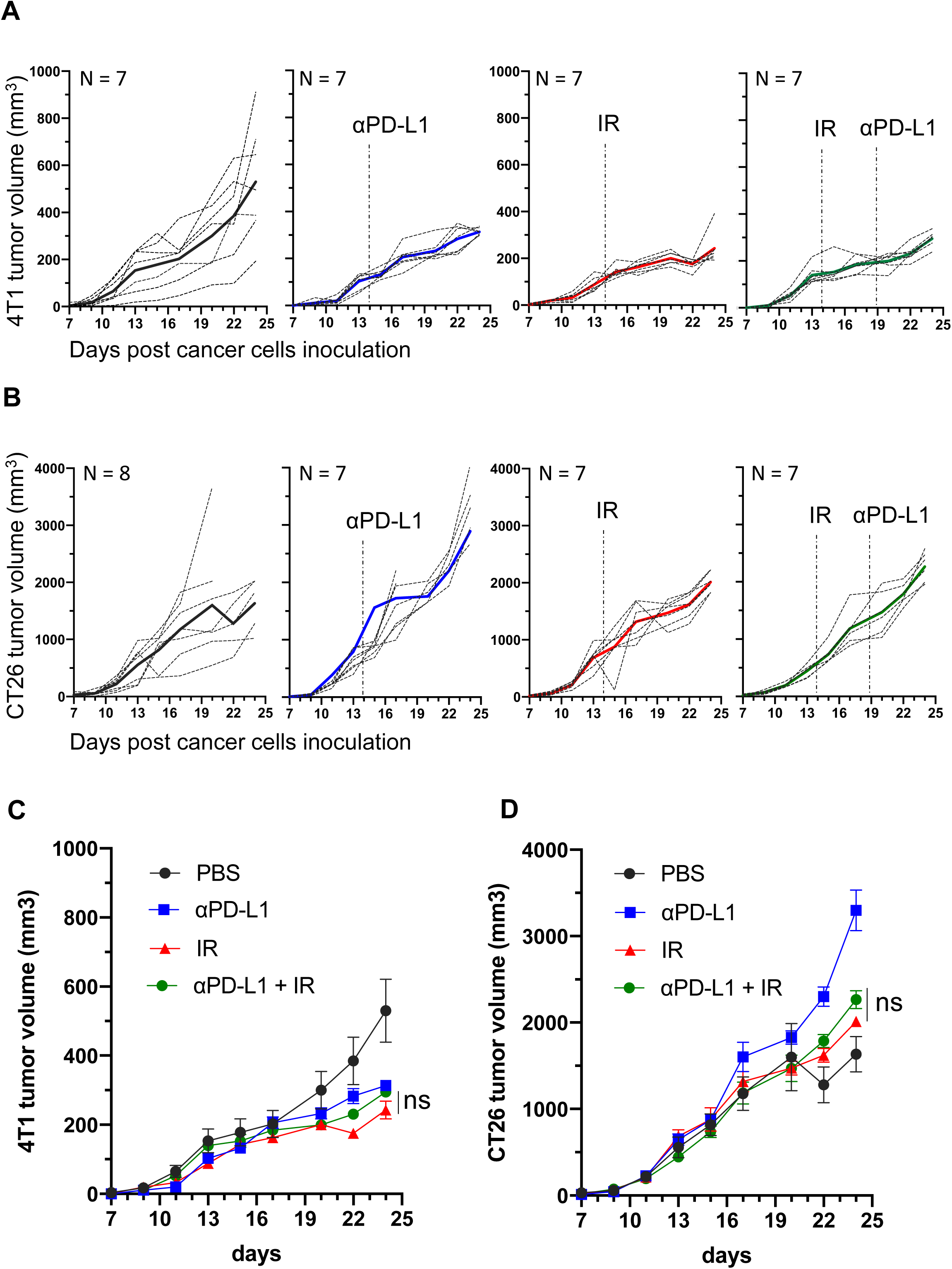
Radiation and anti-PD-L1 are ineffective in immunodeficient NSG mice. Individual growth profiles for 4T1 (**A**) and CT26 (**B**) tumors treated with PBS, anti-PD-L1, 10 Gy, or combination treatment with anti-PD-L1 injected 5 days after 10 Gy. Mean 4T1 **(C)** and CT26 **(D)** tumor growth profiles of the four treatment groups. ns p>0.05, n=7-8.

**Supplemental Figure 3.**
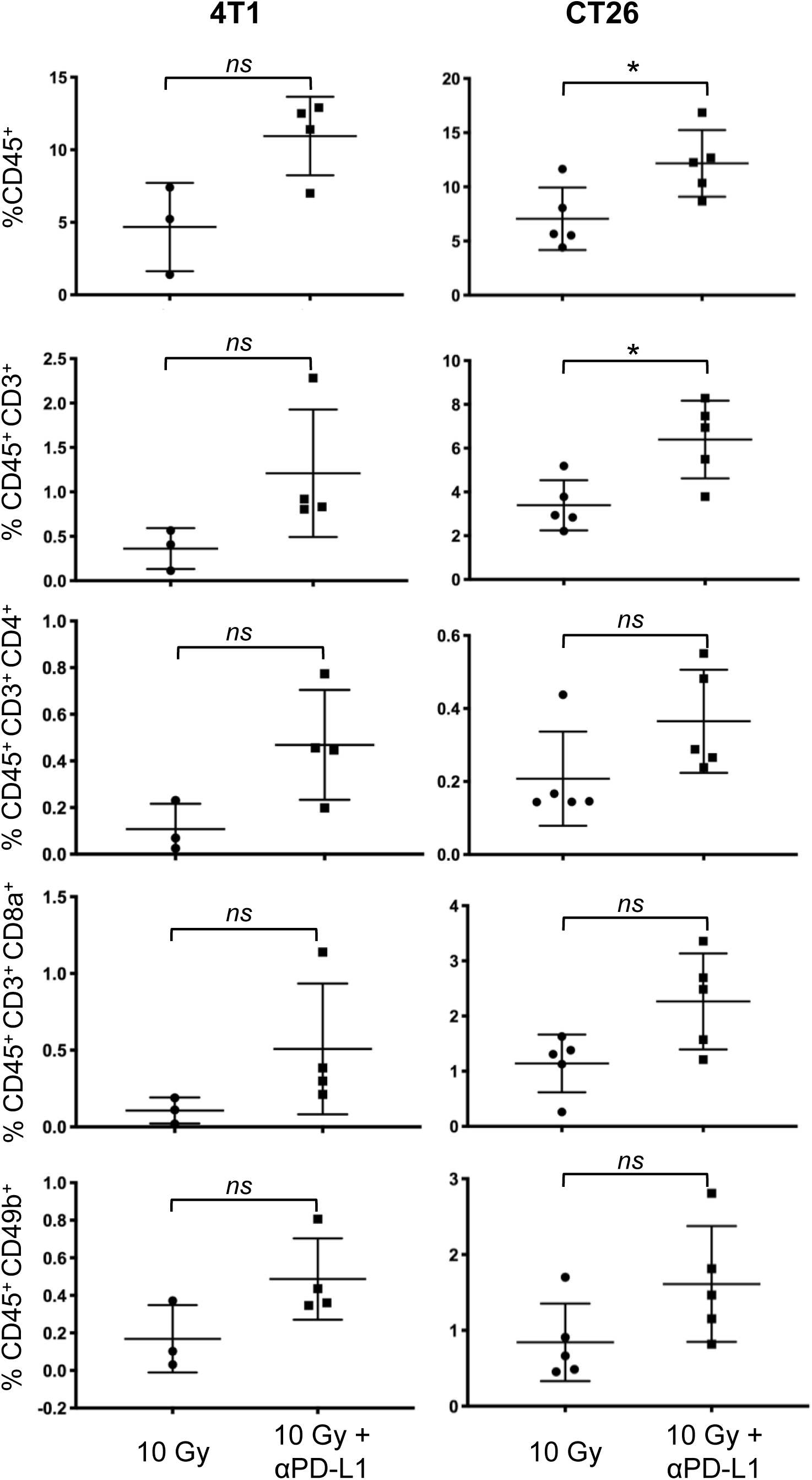
TILs are recruited to tumors by PD-L1 antibody treatment at 5 days post-IR. Flow cytometry was used to quantify the percentage of TILs and key TIL subsets in 4T1 and CT26 tumors treated with 10 Gy alone or with anti-PD-L1 after 5 days. Percentage of total viable cells in samples representing CD45^+^ infiltrating cells, CD45^+^ CD3^+^ T cells, CD45^+^ CD3^+^ CD4^+^ helper T cells, CD45^+^ CD3^+^ CD8^+^ cytotoxic T cells, and CD45^+^ CD49b^+^ natural killer cells. Mean percentage ± SEM, n=3-4 tumors per group, * p<0.05, ns p>0.05.

**Supplemental Figure 4.**
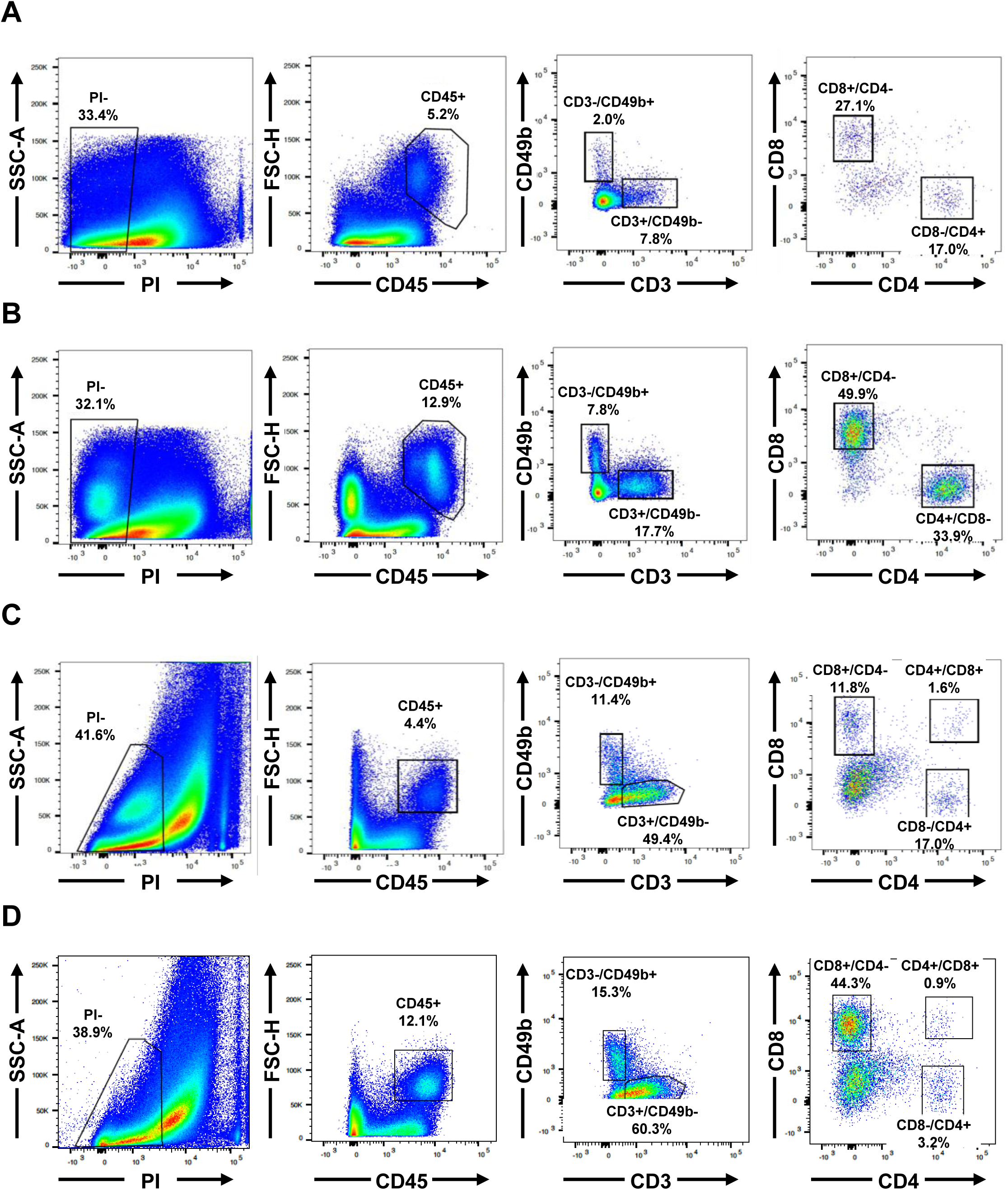
Representative flow cytometry data and gating used in Supplementary Figure 3 for 4T1 tumor treated with 10 Gy (**A**) or 10 Gy and anti-PD-L1 (**B**) and CT26 tumor treated with 10 Gy (**C**) or 10 Gy and anti-PD-L1 (**D**).

## ACKNOWLEDGMENTS

The authors thank current and former colleagues in the Kron and Lee laboratories for advice and support.

## DECLARATIONS

### Ethics Approval and Consent to Participate

Human subjects were not used in this study.

### Consent for Publication

Not required.

### Availability of Data and Material

Not relevant.

### Authors’ Contributions

S.J.K. and S.S.-Y.L. conceived the project. S.S.-Y.L., J.P., S.A., D.S, and Y.L. designed and performed experiments and analyzed data. S.S.-Y.L. and J.P. wrote the manuscript and S.J.K. reviewed the manuscript.

## REFERENCES

1. Chamoto K, Yaguchi T, Tajima M, Honjo T. Insights from a 30-year journey: function, regulation and therapeutic modulation of PD1. Nature Reviews Immunology 2023;23(10):682–95 doi: 10.1038/s41577-023-00867-9

2. Sun Q, Hong Z, Zhang C, Wang L, Han Z, Ma D. Immune checkpoint therapy for solid tumours: clinical dilemmas and future trends. Signal Transduction and Targeted Therapy 2023;8(1):320 doi: 10.1038/s41392-023-01522-4

3. Davis AA, Patel VG. The role of PD-L1 expression as a predictive biomarker: an analysis of all US Food and Drug Administration (FDA) approvals of immune checkpoint inhibitors. Journal for ImmunoTherapy of Cancer 2019;7(1):278 doi: 10.1186/s40425-019-0768-9

4. Li H, van der Merwe PA, Sivakumar S. Biomarkers of response to PD-1 pathway blockade. British Journal of Cancer 2022;126(12):1663–75 doi: 10.1038/s41416-022-01743-4

5. Gong J, Le TQ, Massarelli E, Hendifar AE, Tuli R. Radiation therapy and PD-1/PD-L1 blockade: the clinical development of an evolving anticancer combination. J Immunother Cancer 2018;6(1):46 doi: 10.1186/s40425-018-0361-7

6. Wang NH, Lei Z, Yang HN, et al. Radiation-induced PD-L1 expression in tumor and its microenvironment facilitates cancer-immune escape: a narrative review. Ann Transl Med 2022;10(24):1406 doi: 10.21037/atm-22-6049

7. Gilad Y, Eliaz Y, Yu Y, Han SJ, O’Malley BW, Lonard DM. Drug-induced PD-L1 expression and cell stress response in breast cancer cells can be balanced by drug combination. Scientific Reports 2019;9(1):15099 doi: 10.1038/s41598-019-51537-7

8. Zens P, Bello C, Scherz A, et al. The effect of neoadjuvant therapy on PD-L1 expression and CD8+lymphocyte density in non-small cell lung cancer. Modern Pathology 2022;35(12):1848–59 doi: 10.1038/s41379-022-01139-y

9. Yamazaki T, Vanpouille-Box C, Demaria S, Galluzzi L. Immunogenic Cell Death Driven by Radiation-Impact on the Tumor Microenvironment. Cancer Treat Res 2020;180:281–96 doi: 10.1007/978-3-030-38862-1_10

10. Galluzzi L, Aryankalayil MJ, Coleman CN, Formenti SC. Emerging evidence for adapting radiotherapy to immunotherapy. Nature Reviews Clinical Oncology 2023;20(8):543–57 doi: 10.1038/s41571-023-00782-x

11. Dovedi SJ, Adlard AL, Lipowska-Bhalla G, et al. Acquired resistance to fractionated radiotherapy can be overcome by concurrent PD-L1 blockade. Cancer Res 2014;74(19):5458–68 doi: 10.1158/0008-5472.CAN-14-1258

12. Deng L, Liang H, Burnette B, et al. Irradiation and anti-PD-L1 treatment synergistically promote antitumor immunity in mice. J Clin Invest 2014;124(2):687–95 doi: 10.1172/JCI67313

13. Pointer KB, Pitroda SP, Weichselbaum RR. Radiotherapy and immunotherapy: open questions and future strategies. Trends Cancer 2022;8(1):9–20 doi: 10.1016/j.trecan.2021.10.003

14. Rajeev-Kumar G, Pitroda SP. Synergizing radiotherapy and immunotherapy: Current challenges and strategies for optimization. Neoplasia 2023;36:100867 doi: 10.1016/j.neo.2022.100867

15. Potchen EJ, Kinzie J, Curtis C, Siegel BA, Studer RK. Effect of irradiation on tumor microvascular permeability to macromolecules. Cancer 1972;30(3):639–43 doi: 10.1002/1097-0142(197209)30:3<639::aid-cncr2820300308>3.0.co;2-3

16. Yamazaki T, Young KH. Effects of radiation on tumor vasculature. Mol Carcinog 2022;61(2):165–72 doi: 10.1002/mc.23360

17. Appelbe OK, Zhang Q, Pelizzari CA, Weichselbaum RR, Kron SJ. Image-Guided Radiotherapy Targets Macromolecules through Altering the Tumor Microenvironment. Mol Pharm 2016;13(10):3457–67 doi: 10.1021/acs.molpharmaceut.6b00465

18. Lee SS, Bindokas VP, Kron SJ. Multiplex Three-Dimensional Mapping of Macromolecular Drug Distribution in the Tumor Microenvironment. Mol Cancer Ther 2019;18(1):213–26 doi: 10.1158/1535-7163.Mct-18-0554

19. Liu Y, Cheng Y, Xu Y, et al. Increased expression of programmed cell death protein 1 on NK cells inhibits NK-cell-mediated anti-tumor function and indicates poor prognosis in digestive cancers. Oncogene 2017;36(44):6143–53 doi: 10.1038/onc.2017.209

20. Oh SA, Wu D-C, Cheung J, et al. PD-L1 expression by dendritic cells is a key regulator of T-cell immunity in cancer. Nature Cancer 2020;1(7):681–91 doi: 10.1038/s43018-020-0075-x

21. Monjazeb AM, Schalper KA, Villarroel-Espindola F, Nguyen A, Shiao SL, Young K. Effects of Radiation on the Tumor Microenvironment. Semin Radiat Oncol 2020;30(2):145–57 doi: 10.1016/j.semradonc.2019.12.004

22. Passelli K, Repáraz D, Kinj R, Herrera FG. Strategies for overcoming tumour resistance to immunotherapy: harnessing the power of radiation therapy. Br J Radiol 2024;97(1160):1378–90 doi: 10.1093/bjr/tqae100

